# Chloroplast SRP43 subunit Prevents Aggregation of Proteins

**DOI:** 10.1101/2019.12.24.888255

**Authors:** Mercede Furr, Patience Okoto, Mahmoud Moradi, Colin Heyes, Ralph Henry, Thallapuranam Krishnaswamy Suresh Kumar

## Abstract

Integration of light-harvesting chlorophyll binding proteins into the thylakoid membrane requires a specific chaperone, being the cpSRP43 subunit, of the signal recognition particle pathway in chloroplasts. cpSRP43, unique to the chloroplast, is responsible for transport of LHCPs through the stroma as well as assisting in the correct folding, assembly and disaggregation of these proteins for the acquisition of light energy. cpSRP43 is a highly flexible, multidomain protein capable of binding distinct partners in the cpSRP pathway. cpSRP43 is an irreplaceable component, necessary for the accurate and successful integration of LHCPs. It can act as a disaggregase without any input of external energy. Its action is based on the ability to associate with variable regions of different proteins owing to the domains and flexibility within its distinctive structure. Understanding the unique capabilities of cpSRP43 in the chloroplast begs the question of its usefulness outside of the plant cell, as well as its yet unknown roles still within the plant cell. Although the capabilities of cpSRP43 as a hub protein, adept to binding many unknown partners, has been alluded to in other works, it has yet to be thoroughly investigated. In this study we discover that cpSRP43 can act as a generic chaperone for proteins other than LHCP/not native to the chloroplast. The high thermal stability of cpSRP43 has been demonstrated in the previous chapter by its ability to retain its secondary structure as well as withstand aggregation upon heating and cooling cycles as confirmed by absorbance, intrinsic tryptophan fluorescence and far UV circular dichroism spectroscopy. This property gives cpSRP43 the basis to act as a generic chaperone and provide protection like that of typical heat shock proteins. Carbonic anhydrase, Concanavalin A and hFGF1 (acidic human fibroblast growth factor), were selected as candidates for chaperoning activity by cpSRP43. In all three cases, heat-induced aggregation of the candidate protein was either eliminated or significantly reduced in the presence of cpSRP43. In the case of hFGF1, the bioactivity was preserved after heat-treatment in the presence of cpSRP43. We have proposed a mechanism by which cpSRP43 is able to execute this action however further investigation is warranted to determine the exact mechanism(s) which may vary dependent on the target protein.

## Introduction

The chloroplast signal recognition particle (cpSRP) executes the translocation of Light-harvesting Chlorophyll Binding Proteins (LHCPs) through the chloroplast stroma and assists, in coordination with the cpSRP receptor (FtsY) and the integral membrane protein ALB3, with their integration into the thylakoid membrane (1). The cpSRP consists of the conserved 54-kDa subunit (cpSRP54) and a unique 43-kDa subunit (cpSRP43) (2, 3). cpSRP and LHCP form a soluble “transit complex” thereby enabling the LHCPs to traverse the stroma without becoming involved in inappropriate interactions (1, 4). cpSRP43 is required to prevent aggregation of the hydrophobic LHCPs during their posttranslational targeting to the thylakoid membrane (5). cpSRP43 has been structural identified as having three chromodomains (CD1-3) and four ankyrin repeats (Ank1-4) located between CD1 and CD2 (4, 6). The crystal structure of cpSRP43 reveals two hydrophobic grooves which are disjointed by a positive ridge on one side and a highly negatively charged surface on the other side (7). The interaction between cpSRP and LHCP is achievable, in part, by the binding between the ANK regions of cpSRP43 and the L18 domain of LHCP, an 18-amino acid, hydrophilic peptide which connects the second and third trans membrane domains (TM2, TM3) (6, 8). The chromodomains located on the c-terminal of cpSRP43 allow for its ability to bind the methionine-rich domain of cpSRP54 lending to the conformation of the transit complex (6). In addition, cpSRP43 binds to portions of the integral membrane protein ALB3 to initiate and expedite docking at the receptor for integration. The unique architecture and multi-domains which make up cpSRP43 make it an excellent framework upon which protein interactions can occur.

Investigation into the structure and chaperone cycle of cpSRP43 suggests that it is highly specific for chaperoning LHCPs (9). The aggregation of LHCPs was monitored in the presence of each cpSRP pathway component and cpSRP43 was shown to be the exclusive component for preventing the aggregation of LHCPs (5). cpSRP43 does not require ATPase activity as is the case for the Hsp104/ClpB family of chaperones, however its ability to dissolve aggregates of LHCPs is comparable to that family of disaggregases (9–12). Studies have also revealed that Hsp70, Hsp60 and trigger factor (TF), a bacterial chaperone involved with hydrophobic regions of proteins, cannot substitute for cpSRP43 indicating the specificity of cpSRP43 to LHCPs (5, 13, 14). Hydrophobic interactions between the trans-membrane regions of LHCP and cpSRP43 are also considered to contribute largely to the ability of cpSRP43 to successfully chaperone and convoy the LHCPs to their destination. The contribution of hydrophobic interactions has been asserted based on studies in which binding was not lost under high salt conditions, a 60-fold higher binding affinity between the full length LHCP versus the L18 peptide alone and the involvement of TM3 for efficient binding (5, 15, 16).

Proteins which act as molecular chaperones are part of a diversified classification due to their scope of specific functions (17). Proteins in this group maintain crucial roles in folding and assembly, refolding, translocation and preventing aggregation (17). In addition, chaperones assist under stress and disease conditions. The biogenesis of an abundant amount of membrane proteins typically takes place in conjunction with their insertion into the membrane, however, in posttranslational translocation of membrane proteins, the requirement of effective chaperones is imperative. Molecular chaperones which can prevent, and/or reverse protein aggregation provide a significant contribution in the maintenance of cellular stability, proper function and prevention of disease states. Molecular chaperones known as “disaggregases” are capable of reversing protein aggregation with the aid of external energy provided by ATP hydrolysis as well as other co-chaperones (14, 18, 19). cpSRP43 is one of the first proteins to be described as an ATP-independent disaggregase. cpSRP43 provides protection to the most abundant protein on earth, giving it a highly specific and critical function. To investigate how cpSRP43 can recognize and reconstruct LHCP aggregates, the characteristics and structure of LHCP aggregates was elucidated (11, 20). LHCP aggregates are highly stable, insoluble, resistant to detergents or reversal by dilution (11). The L18 peptide and TM3 are shown to be solvent exposed in the aggregate formations which presents as the recognition site for cpSRP43 to first interact with the LHCP aggregate and begin the disassembly and reorganization of each LHCP protein (11). Recent studies using NMR and smFRET have also shown cpSRP43 to have high conformational flexibility (21, 22).

cpSRP43 is an indispensable component of the cpSRP pathway, involved in interactions with multiple partners leading to the successful translocation, disaggregation, refolding and integration of LHCPs. The ability of cpSRP43 to disassemble and remodel LHCP aggregates relies on its binding interactions with its substrate in place of any external energy source and its flexibility demonstrates its effectiveness and potential adaptability as a generic chaperone. Although LHCPs specifically require cpSRP43 as a chaperone, it stands to reason that cpSRP43 may also serve as a general chaperone which prevents aggregation and assists in the refolding of other proteins. The distinctive heat stability of cpSRP43 was outlined in the previous chapter. In this study, we demonstrate its ability to assist in refolding and prevention of heat-induced aggregation of three proteins which cpSRP43 from *Arabidopsis Thaliana* does not ordinarily associate with by using Carbonic anhydrase from bovine erythrocytes, Concanavalin A from *Canavalia ensiformis* (Jack Bean) and, hFGF1 as substrates.

## Materials and Methods

### Expression and Purification of Recombinant cpSRP43

BL-21-star cells were transformed with the expression vector pGEX T2 encoding the sequence of cpSRP43 grown at 37°C lysogeny broth containing 10 μg/ml ampicillin. Protein expression was induced at an optical density of 0.8 measured at A600 with1mM isopropyl- Dthiogalactoside. The cells were incubated for an additional 3.5 hours followed by harvesting by centrifugation. Cell pellets were resuspended and lysed by sonication for 25 cycles with 10 second on/off pulses using 10 W of output power. GST-cpSRP43 was bound to glutathione Sepharose then washed thoroughly with equilibration buffer (2.7mM KCl, 1.8mM KH_2_PO_4_, 15mM Na_2_HPO_4_, 137mM NaCl, pH 7.2). GST-cpSRP43 was eluted with 10 mM L-Glutathione and exchanged into cleavage buffer (50 mM Tris-HCl, 150 mM NaCl, 1 mM EDTA, 1 mM dithiothreitol, pH 7.0). Overnight cleavage was setup, in solution, at 4°C on a rocker with 10 units of PreScission Protease per liter of original cells for 16 hours. The cleavage product was again put onto glutathione for separation of the GST tag and the cleaved cpSRP43 was further purified by size exclusion chromatography (SEC) as seen in Fig. 1 below.

**Fig.1:**
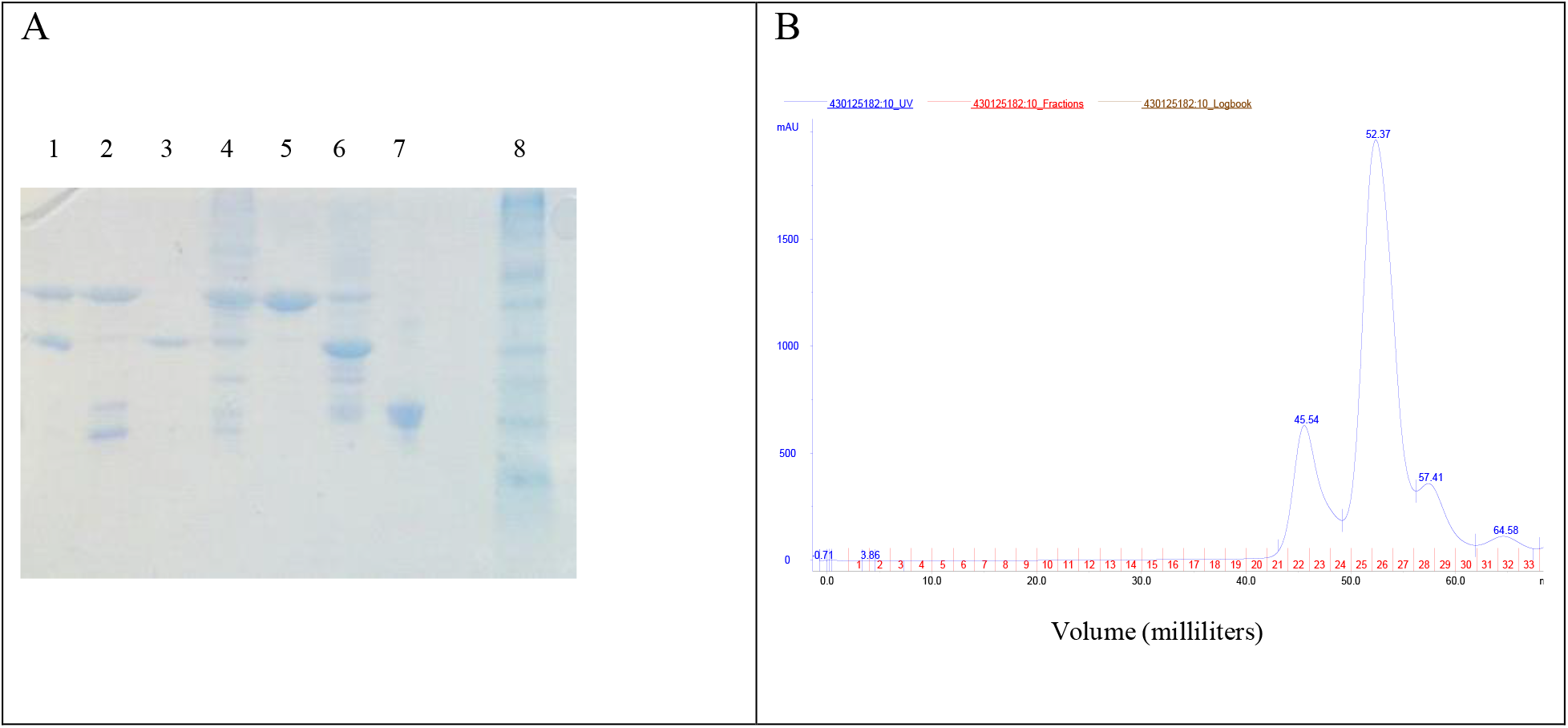
Purification of cpSRP43, SDS-PAGE (Panel-A) and the SEC profile (Panel-B) of cpSRP43 purification. Panel-B: Lane 1= Cleavage product containing cpSRP43 and GST, Lane 2= cleaved cpSRP43 eluted in the unbound fraction from GSH column, Lane 3= GST tag eluted with 10 mM L-Glutathione, Lane 4= Peak 1 from SEC eluted at 45 milliliters containing cpSRP43 and contaminants, Lane 5= Peak 2 from SEC eluted at 52 milliliters containing cpSRP43, Lane 6= Peak 3 from SEC eluted at 57 milliliters containing cpSRP43, GST and contaminants Lane 7= Peak 4 eluted from SEC containing low molecular weight contaminants.

### Expression and Purification of hFGF1

hFGF1 was overexpressed in BL-21(DE3) cells and grown to an Optical Density of 0.6–1.0 at Abs_600_ and incubated with 1 mM isopropyl b-Dthiogalactoside for 3.5 hours. Cells were harvested, resuspended and sonicated. The supernatant was separated from the cell debris using ultra centrifugation. hFGF1 was purified on a heparin sepharose column using a stepwise salt gradient in 10mM sodium phosphate buffer containing 25mM (NH_4_)_2_SO_4_ pH 7.2 and purity was analyzed using 15% SDS-PAGE (Fig.2). Pure hFGF1 elutes at a concentration of 1500 mM NaCl.

**Fig.2:**
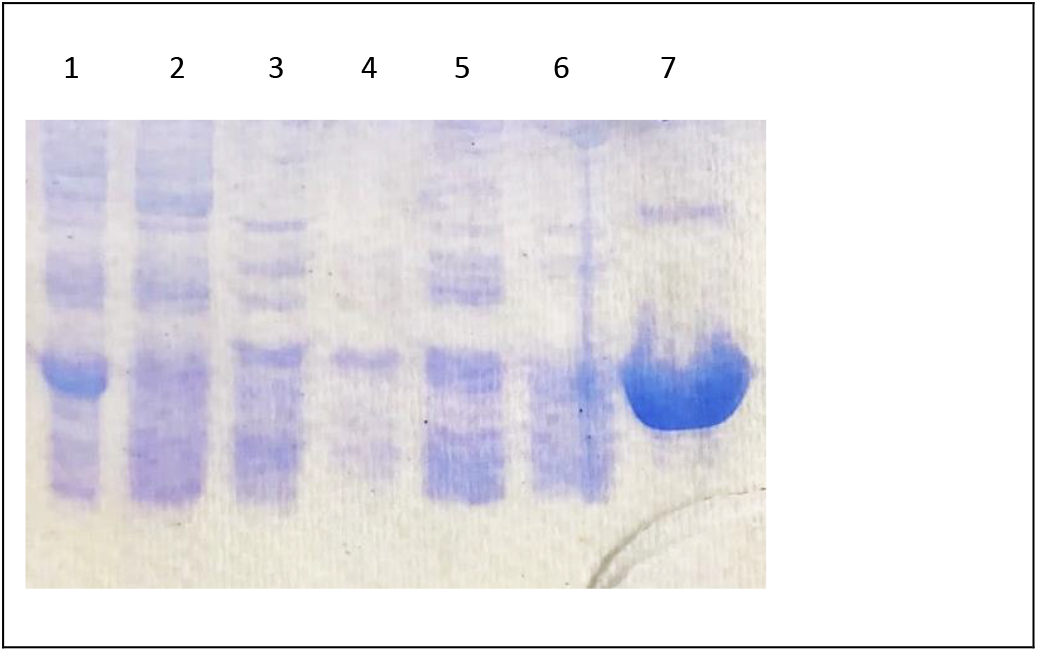
Purification of wtFGF1 on Heparin Sepharose column. Lane 1= Supernatant, Lane 2= Unbound fraction, Lane 3= 100mM NaCl elution, Lane 4= 300 mM NaCl elution, Lane 5= 500 mM NaCl elution, Lane 6= 800 mM NaCl elution, Lane 7= 1500 mM NaCl elution.

**Carbonic Anhydrase** from bovine erythrocytes and **Concanavalin A** from *Canavalia ensiformis* (Jack Bean) Type V were purchased commercially (Sigma).

### Circular Dichroism

Circular dichroism (CD) measurements were performed on a Jasco J-1500 CD spectrometer equipped with a variable temperature cell holder. Conformational changes in the secondary structure of cpSRP43 were monitored in the far-UV region between 190 to 250 nm with a protein concentration of 0.3mg/ml in a quartz cuvette with a pathlength of 1 mm. The scanning speed, band width and data pitch were set to 50 nm/min, 1.00 nm and 0.1nm, respectively. Three scans were taken (within a 1000 HT voltage range) and averaged to obtain the CD spectra. The thermal denaturation/renaturation scans were recorded from 25°C to 90°C at 5°C increments.

### Size-Exclusion Chromatography (SEC)

cpSRP43, hFGF1 and cpSRP43/hFGF1 proteins for the mixture samples were over-expressed and purified as described above. In the heat-treated and non-heat-treated samples, concentrations of hFGF1 and cpSRP43 used were 20 and 25 μM, respectively. The heat-treated and non-heat-treated samples were loaded to a Superdex 75 16/600 column (GE Healthcare, Pittsburgh, PA) equilibrated in 2.7mM KCl, 1.8mM KH_2_PO_4_, 15mM Na_2_HPO_4_, 137mM NaCl, pH 7.2 on an AKTA FPLC and ran at a flow rate of 1 milliliter per minute. The elution volume was monitored by absorbance at 280 nm. Fractions were collected and analyzed by SDS-PAGE.

### Refolding of Carbonic Anhydrase in the presence and absence of cpSRP43 as monitored by turbidity measurements

All readings were obtained using an Agilent UV-VIS Spectrophotometer. Absorbance readings were recorded at 280 nm and 350 nm in order to track concentration and aggregation, respectively. Carbonic anhydrase was denatured and reduced in 8 M Urea with 1 mM BME and added to refolding buffer (10 mM phosphate, 100 mM NaCl, 0.1 mM GSH and 1 mM G-S-S-G at pH 7.2) starting at 0.1 mg/ml and increasing by 0.1 mg/ml for each reading. A total of ten readings were acquired for each experiment and the experiments were repeated three times each for denatured and reduced carbonic anhydrase increasing in refolding buffer in the presence and absence of cpSRP43. A constant concentration of 0.2 mg/ml (6 μM) of cpSRP43 was present in the refolding experiments carried out in the presence of cpSRP43.

### Temperature-Dependent Aggregation Inhibition Experiments

All absorbance readings were recorded at 280 and 350 nm wavelengths using an Agilent UV-VIS Spectrophotometer. The increasing temperatures were maintained using a variable temperature water-bath. Carbonic Anhydrase, Concanavalin A and hFGF1 were each heated at 5°C increments in the presence and absence of cpSRP43. All samples were held at each temperature increment for 5 minutes prior to measurement. The concentration of cpSRP43 was held constant for each substrate protein in the presence of cpSRP43 experiments. The concentrations of the substrate proteins were increasing for each reading in the presence and absence of cpSRP43.

### Bioactivity Assays

3T3 fibroblast cells obtained from ATCC (Manassas, VA) were cultured in media consisting of DMEM supplemented with 10% bovine calf serum. Cells were grown and were incubated overnight at 37 °C. The bioactivity of FGF1 was determined by quantifying the cell number increase after being incubated with FGF1 post heat treatment at 60°C and FGF1 in the presence of cpSRP43 post heat treatment at 60°C. Starved 3T3 fibroblasts were collected and seeded in a well plate at a seeding density of 10,000 cells/well. The bioactivity assays were performed five times under the same condition. 3T3 cell proliferation was assessed by the Cell Titer-Glo (Promega, Madison, WI) cell proliferation assay after 24 hours.

## Results and Discussion

### cpSRP43 possesses high thermal stability

cpSRP43 shows a circular dichroism spectrum with negative bands at 208 and 222 nm which indicate that it has a predominate, typical α helical secondary structure. An intrinsic fluorescence emission at 341 nm indicates that the protein is in its native conformation. Thermal denaturation of cpSRP43 was probed by circular dichroism from 25-90°C at 5°C increments and retained its secondary structure both upon heating and cooling cycles as shown in the previous chapter. cpSRP43 is heat stable up to 90°C and can retain its secondary structure during heating. In addition, cpSRP43 was heated at five-degree increments from 25 to 90°C and the absorbance at 350 nm was recorded after the protein was held for five minutes at each temperature. cpSRP43 did not show visible aggregation or aggregation reflected by absorbance readings up to 90°C (Fig.3B). Subsequently, ultracentrifugation of the post heat treated samples did not indicate protein precipitation upon visual inspection.

**Fig.3:**
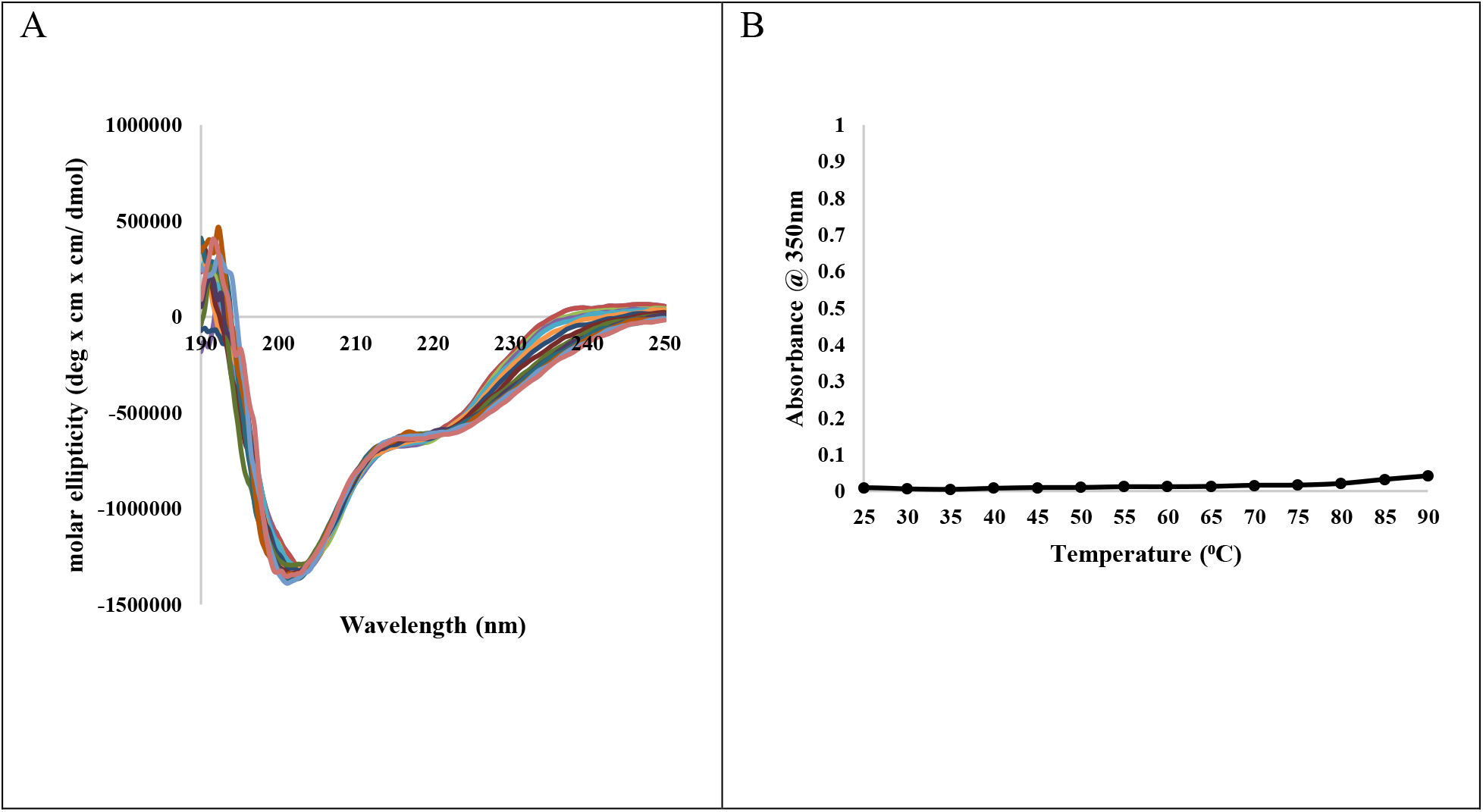
Overlay of Far-UV CD spectra of cpSRP43 upon thermal denaturation (Panel-A). Aggregation profile of cpSRP43 (Panel-B) upon heat treatment as probed by absorbance at 350 nm.

### cpSRP43 assists in the refolding of denatured and reduced Carbonic Anhydrase

Carbonic anhydrase (CA), found in all kingdoms of life, catalyzes the reversible conversion of carbon dioxide and water to bicarbonate and also plays a role in ion transport, pH regulation and water and electrolyte balance (23, 24). CA II deficiency is responsible for osteopetrosis, an inherited syndrome in humans which is characterized with renal tubular acidosis and brain calcification (25–28). CA has been studied more closely over the past decade for its role in biological systems pertaining to bone resorption, renal acidification, and for its suggested importance in normal brain development (25, 28). CA was used in this study to illustrate the ability of cpSRP43 to aid in the refolding of a protein other than LHCP and although CA is naturally abundant in chloroplasts, the CA used here originated from bovine erythrocytes. A mixture of reduced and oxidized glutathione was incorporated in the refolding buffer for its usefulness in facilitating and/or accelerating disulfide bond formation once destroyed by solubilization in a denaturant(29–31). A stock solution of CA was denatured and reduced in 8 M Urea containing 1 mM BME. A Urea buffer was prepared containing (10 mM phosphate, 100 mM NaCl, 8M Urea and 1 mM BME at pH 7.2) and a refolding buffer was prepared containing (10 mM phosphate, 100 mM NaCl, 0.1 mM GSH and 1 mM G-S-S-G at pH 7.2) Increasing concentrations of CA were added (0.1mg/ml increments) to refolding buffer in the presence and absence of cpSRP43 (Fig 4). In refolding buffer that did not contain cpSRP43, the CA began to aggregate at a concentration of 0.2 mg/ml. In refolding buffer containing cpSRP43 (constant concentration of 0.2 mg/ml), the CA did not show any aggregation as probed by absorbance readings taken at 350 nm until the CA reached a concentration of 0.5 mg/ml and did not show visible aggregation until the concentration reached 0.6 mg/ml (Fig.4). It appears that without the assistance of cpSRP43 the CA was aggregating almost immediately, and the aggregates were visible at a concentration of 0.3 mg/ml. The refolding buffer composition alone was not adequate to reduce the rate of aggregation of CA however the presence of cpSRP43 in the refolding buffer did postpone aggregate formation up to three times that of the refolding buffer alone. This result indicates that cpSRP43 can assist in the refolding of CA, being a concentration dependent action.

**Fig. 4:**
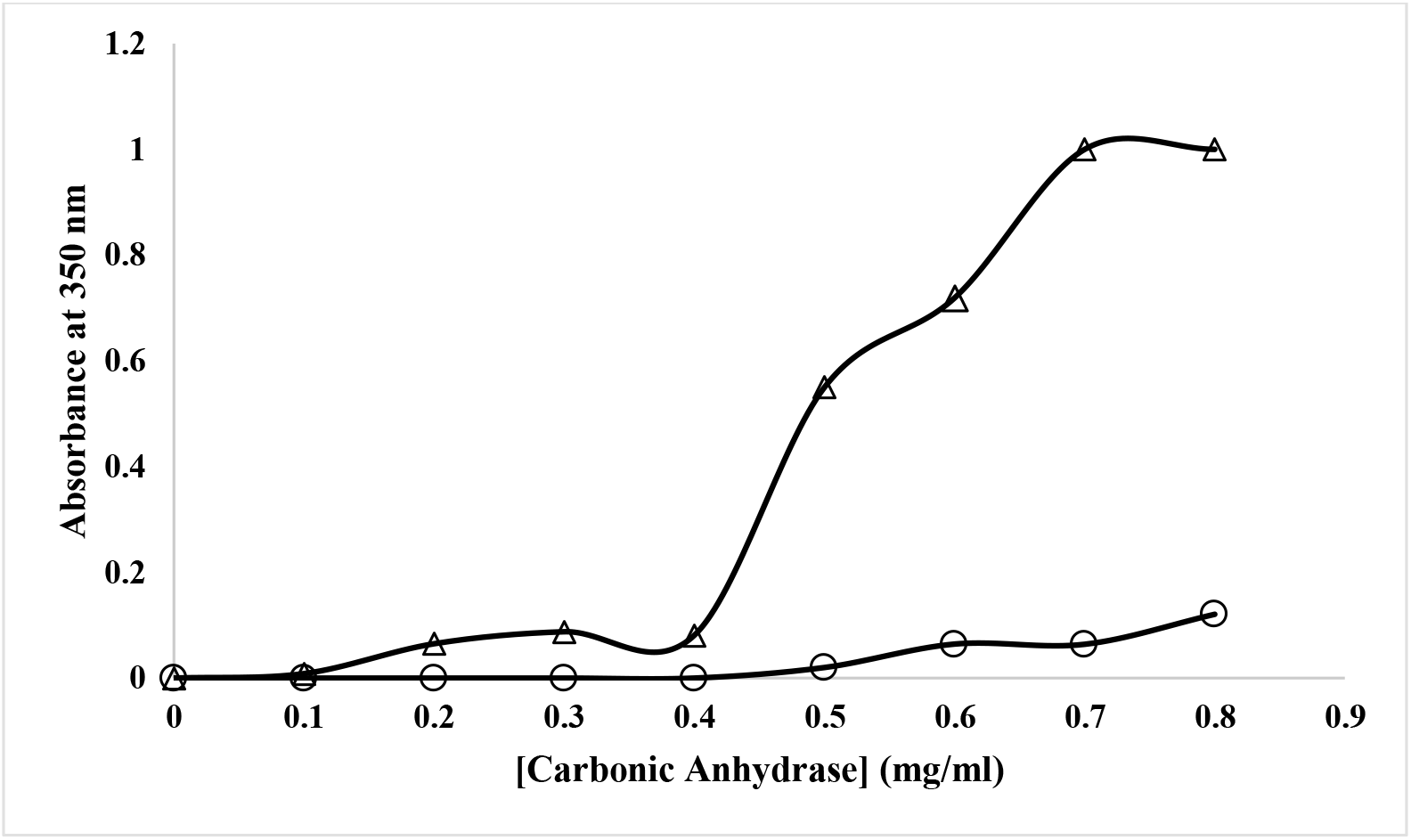
Refolding of carbonic anhydrase in the presence (o) and absence (Δ) of cpSRP43

### cpSRP43 prevents heat-induced aggregation of Carbonic Anhydrase and Concanavalin A

The chaperone ability of cpSRP43 was investigated by monitoring the heat-induced aggregation of different target proteins in the absence and presence of cpSRP43. A preliminary heat treatment experiment done using CA alone determined that CA was heat stable up to 60°C therefore the experiment was carried out from a starting temperature of 40°C. Increasing concentrations of carbonic anhydrase, in the presence and absence of cpSRP43, were heated from 40-90°C at 5°C increments and held for 5 minutes at each temperature before readings were taken at an absorbance of 350 nm. Each concentration of CA was added to eleven sample tubes (1 for each temperature) and made up with buffer in the control experiments and buffer containing 25 μM of cpSRP43 in the presence of cpSRP43 experiments prior to heating. The experiment was repeated at increasing concentrations of CA in the absence and presence of a constant concentration of cpSRP43. At the micromolar ratio in which cpSRP43 was in the most excess to CA (0.2: 1, Fig. 5A(a)), its presence was able to provide the best degree of protection. In the control experiment with CA alone it became completely aggregated by 70°C and at the same temperature in the presence of excess cpSRP43 there was still some aggregation present, but it was nine times less the amount of aggregation as in the control experiment. At equivalent ratios it appears that protection from heat-induced aggregation was not provided by cpSRP43 suggesting that the event in which cpSRP43 is acting as a chaperone for CA is concentration dependent (Fig.5A(c)). A similar tread was seen in the same experiments in which Concanavalin A (ConA) was used as the target protein for chaperoning activity provided by cpSRP43, however a better rate of protection was detected between Con A and cpSRP43 in comparison (Fig.5B).

**Fig. 5:**
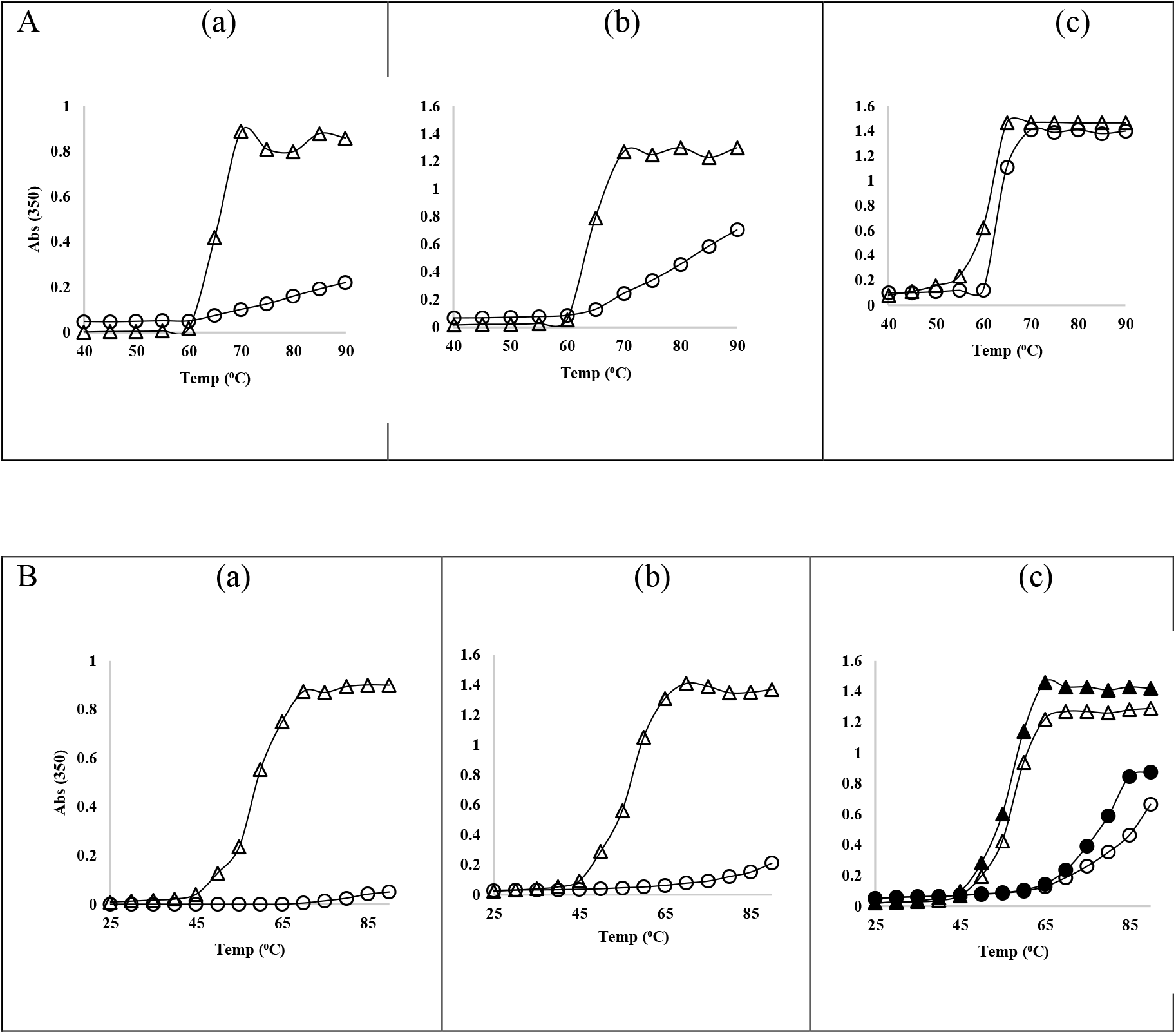
(Panel-A) Heat-induced aggregation profile of Carbonic anhydrase in the presence (o) and absence (Δ) of cpSRP43. (Panel-B) Heat-induced aggregation profile of Concanavalin A. Concentrations are reported in micromolar ratios: Panel-A(a)(0.2:1), (b)(0.4:1), (c)(1:1). Panel-B(a)(0.13:1),(b)(0.27:1),(c)(Δ:o)(0.36:1), (c)(▲:●)(0.45:1).

Con A is a lectin (carbohydrate-binding protein) which was initially isolated from the jack bean (*Canavalia ensiformis*) (32). It is a homotetramer at pH 7.0 containing 26.5 kDa subunits which bind to the non-reducing terminal α-D-glucosyl and α-D-manosyl groups (32, 33). Lectins are known to play physiological roles in animals but the roles of plant lectins are less understood (34). Although, Con A derived from plant proteins have proven to be especially useful as a probe in the study of cell surface membrane dynamics and mitosis in lymphocytes (33, 35, 36). In chaperoning activity experimentation between Con A and cpSRP43, increasing concentrations of Con A, in the presence and absence of cpSRP43, were heated from 25-90°C at 5°C increments and held for 5 minutes at each temperature increment before readings were taken at an absorbance 350 nm, the same as described above for CA. Significant protection from heat-induced aggregation was observed in the mixture samples of Con A and cpSRP43 in which the ratio of Con A to cpSRP543 was lower i.e. cpSRP43 was in excess (Fig.5B). In all control experiments of Con A, aggregation began at 45°C and complete aggregation was measured and visually observed at 65°C. At 65°C, at Con A to cpSRP43 ratios of 0.13:1 and 0.27:1 micromolar, heat-induced aggregation of Con A was eliminated entirely and reduced by nearly 100%, respectively (Fig.3B (a) and (b)). In the heat-treatment experiments executed at higher ratios of Con A to cpSRP43 protection against aggregation was not significantly observed (Fig.5B (c)) which again supports the assertion that the chaperoning activity of cpSRP43 *in vitro* is concentration dependent.

### cpSRP43 protects hFGF1 and lysozyme from heat-induced aggregation

Human fibroblast growth factor (hFGF1) was selected to display the chaperoning effects of cpSRP43 on a protein which is not native to the chloroplast. Members of the FGF family are involved in signaling cell growth, differentiation and proliferation in addition to having roles in tissue repair, vascular stability and stress response (37–39). FGFs act by binding specific cell surface receptors (FGFRs) (38, 39). FGF1 is considered a heat-labile protein since it has a T_m_ of approximately 40°C which is close to physiological temperature (40, 41). The process by which the heat-induced aggregation inhibition experiments were executed are the same as previously described for CA and Con A. Absorbance readings of hFGF1 alone show that most of the protein has aggregated at 40°C and hFGF1 is completely aggregated at 45°C. Absorbance readings of cpSRP43 alone at 350 nm do not show any turbidity upon heating from 25°C-90°C as shown previously. Heat-induced aggregation of hFGF1 is eliminated at a micromolar ratio of hFGF1 to cpSRP43 of 0.4:1 for up to 70°C (Fig.6A(a)) and up to 65°C at a ratio of 0.8:1 (Fig.6A(b)). Considering that in the absence of cpSRP43, hFGF1 was completely aggregated 45°C the ability to delay this aggregation for 25 more degrees C is significant. In the previous experiments with CA and Con A, when the molar ratios were equivalent, cpSRP43 did not provide any protection against aggregation. However, in the case of hFGF1, at a ratio of 1.2:1 of hFGF1 to cpSRP43, hFGF1 is protected at 45°C, when it becomes completely aggregated otherwise (Fig.6A(c)). At higher molar ratios, protection is only seen at 40°C which indicates that when the quantity of hFGF1 exceeds that of cpSRP43 there is no reduction in the normal aggregation level of hFGF1 at its Tm (Fig.6A(d) and (e)). The protection which cpSRP43 provides to the heat induced aggregation of hFGF1 is concentration dependent with the best protection provided by cpSRP43 at an excess of cpSRP43. This result is consistent throughout experimentation with all three substrates.

**Fig. 6:**
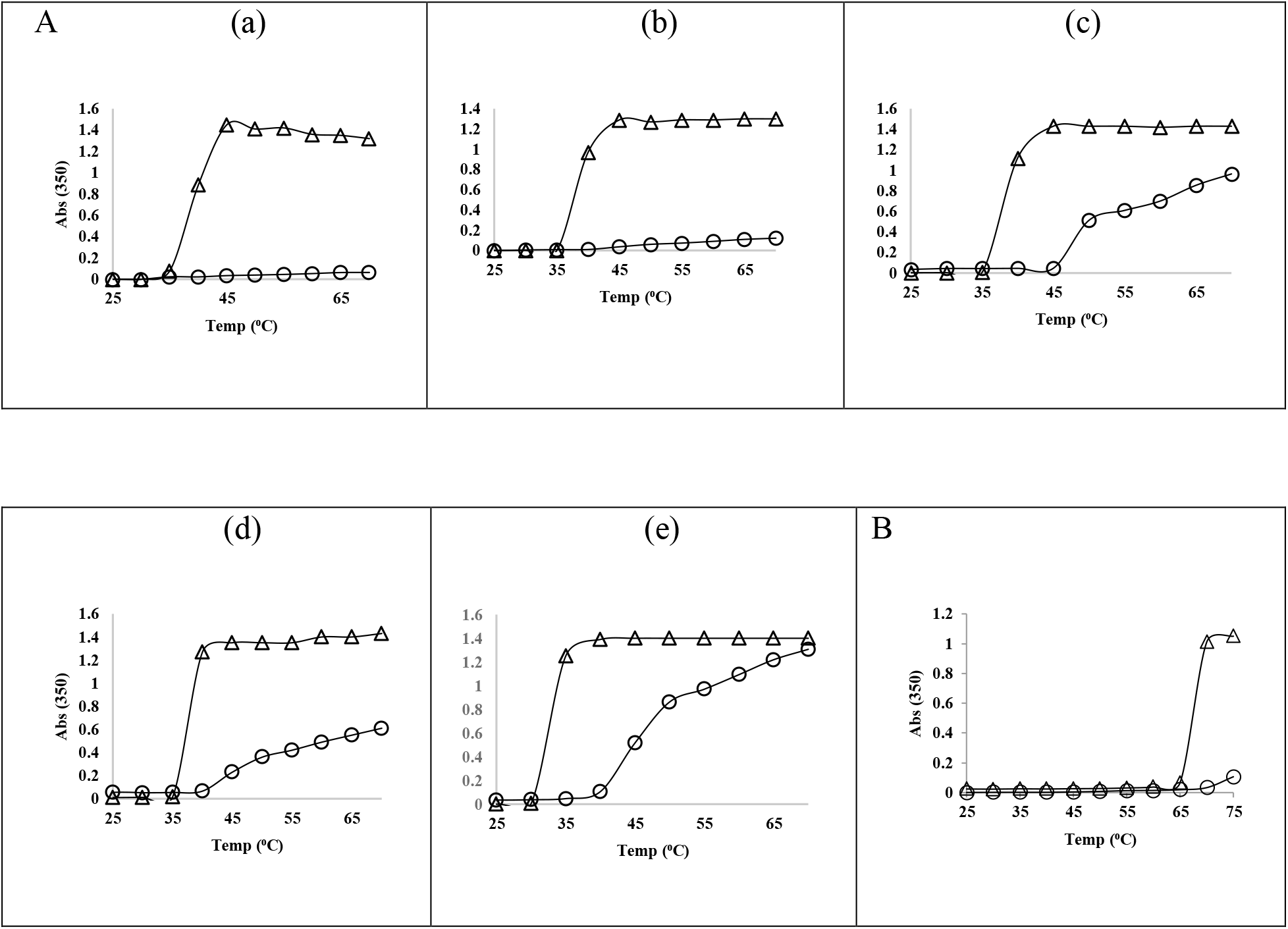
(Panel-A) Heat-induced aggregation profile of hFGF1 in the presence (o) and absence (Δ) of cpSRP43. (Panel-B) Heat-induced aggregation profile of lysozyme in the presence (o) and absence (Δ) of cpSRP43. Concentrations are reported in micromolar ratios: Panel-A(a)(0.4:1), (b)(0.8:1), (c)(1.2:1), (d)(1.6:1), (e)(2.4:1). Panel-B (0.8:1)

In addition to the afore mentioned chaperone activity of cpSRP43 candidate proteins, lysozyme was also investigated. Lysozyme is an enzyme which catalyzes the breakdown of cell walls within certain bacteria as a component of the innate immune response in animals (42, 43). As a result of its antibacterial activity it is present in saliva, tears, mucus and human milk as well as being used in medicine and as a food preservative (44, 45). Lysozyme is already a highly heat-stable protein having a T_m_ of 72°C at pH 7.0 (46). In the control experiment, lysozyme aggregated completely at 70°C (Fig.4B). In the corresponding experiment in which the same concentration of lysozyme was heated in the presence of cpSRP43 at a micromolar ratio of lysozyme to cpSRP43 of 0.8:1, aggregation was eliminated at 70°C was decreased by 90% at 75°C (Fig.4B). Although cpSRP43 was not in great excess in this trial it still provided significant protection against heat-induced aggregation of lysozyme at temperatures at which this enzyme would have been completely aggregated. Testing the biological/enzymatic activity of each of these post heat treated chaperoning candidates would have been ideal. However, hFGF1 was chosen for investigation into its bioactivity post heat treatment in the presence of cpSRP43 and yielded positive results as reported in the following section.

### Post heat-treated hFGF1 (in the presence of cpSRP43) retains its cell proliferation activity

The cell proliferation activity of hFGF1 on NIH 3T3 cells was quantified using post heat treated hFGF1 only and a post heat treated mixture of hFGF1 and cpSRP43 and hFGF1 only as a control. The same experiment was repeated five times. The concentrations of hFGF1 and cpSRP43 were the same in all trials. This was to examine whether or not hFGF1 retained its bioactivity after heating in the presence and absence of cpSRP43 and if so to what extend as compared to non-heat treated hFGF1. Bioactivity assays were preformed on hFGF1 which did not undergoes heat treatment to represent the normal amount of cell proliferation expected. The same concentration of hFGF1 was heated at 50°C in the presence and absence of cpSRP43. Both samples were centriguged at 13,000 rpm for 10 minutes. The supernatant of centrifuged, post heat-treated hFGF1 only sample was carefully removed from the precipitate and tested for bioactivity. The post heat treated mixture of hFGF1 to cpSRP43 used was at a micromolar concentration ratio of 0.4:1 since this concentration which showed the best result in the heat-induced aggregation inhibition experiments. Although the bioactivity of hFGF1 in the post heat-treated mixture did not reach the same levels as non-heat treated hFGF1, the presence of cpSRP43 during heating did lead to significantly better cell proliferation in comparison with the heated hFGF1 sample in the absence of cpSRP43 (Fig.7).

**Fig. 7:**
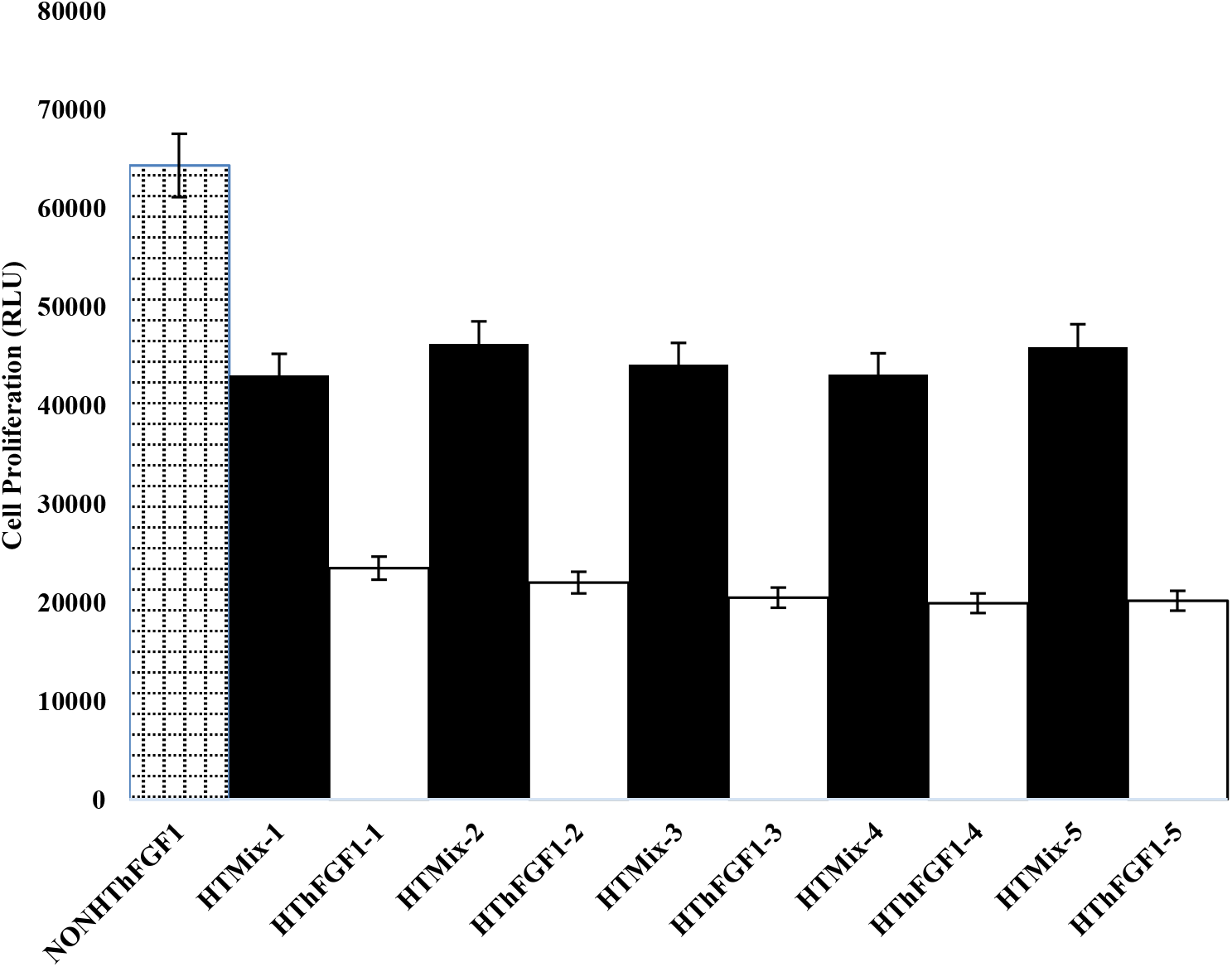
Proliferation of NIH 3T3 cells by heat-treated hFGF1 (HThFGF1-5) only and the heat- treated hFGF1/cpSRP43 mixture (HTMix1-5), repeated five times, samples 1-5. Cell proliferation of non-heat-treated hFGF1 is shown in the far left column of the bar gragh (NONHThFGF1). The standard errors were calculated from triplicate measurements.

Bioactivy of hFGF1 after heat treatment in the presence of cpSRP43 was doubled when compared to the cell proliferation activity of heat treated hFGF1 in the absence of cpSRP43. This is due to the fact that cpSRP43 was able to protect heated hFGF1 from aggregation. A mechanism by which cpSRP43 is able to provide this protection is explored in the dissusion and conclusions section.

### Size exclusion chromatography reveals an interaction between cpSRP43 and hFGF1 under heated conditions which is not present under non-heated conditions

Size exclusion chromatography was performed on heat-treated and non-heat-treated samples of the cpSRP43/hFGF1 mixtures (Fig.8 and 9). The chromatograph of the non-heat-treated mixture contained two well separated peaks at the appropriate elution volumes for cpSRP43 and hFGF1, at 45-52 milliliters and 76 milliliters, respectively (Fig.8A). The corresponding SDS-PAGE revealed a protein band of cpSRP43 and hFGF1 in the separate peaks (Fig.8B). This result suggests that cpSRP43 and hFGF1 do not bind or form associations with each other when combined and incubated at room temperature (20-25°C). The heat-treated SEC profile shows less resolution between the peaks with cpSRP43 eluting at its normal volume (45-52 milliliters) and hFGF1 eluting slightly earlier at 68 milliliters (Fig.9A). The results suggest that, upon heating, cpSRP43 and hFGF1 undergo transient associations although it is not indicative of tight binding.

**Fig. 8:**
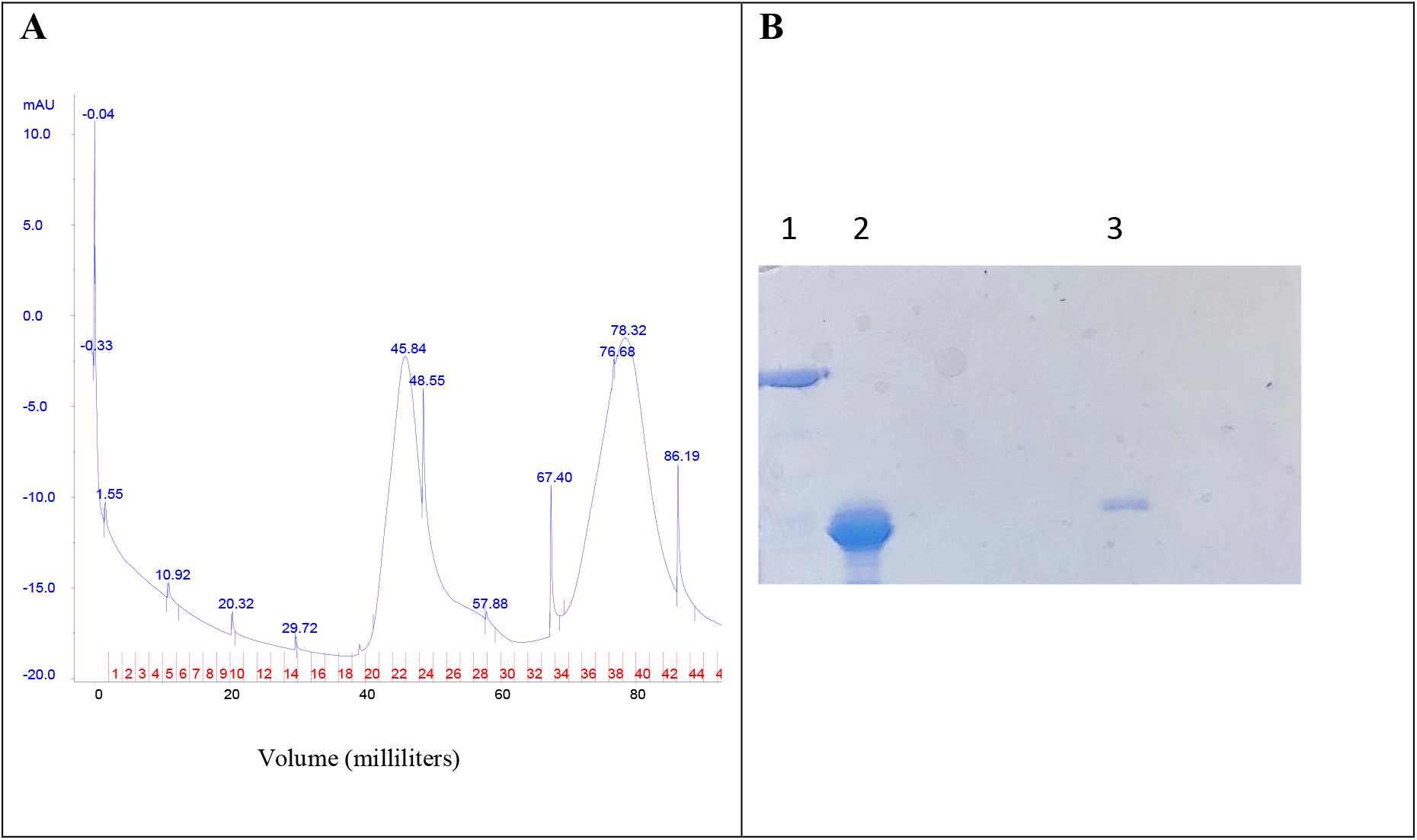
Size-exclusion Chromatography (Panel-A) and SDS-PAGE analysis (Panel-B) of the non-heat-treated mixture of hFGF1 and cpSRP43. Panel-B: Lane 1= Peak 1 collected containing cpSRP43, Lane 2= hFGF1 marker, Lane 3= Peak 2 collected containing hFGF1.

**Fig. 9:**
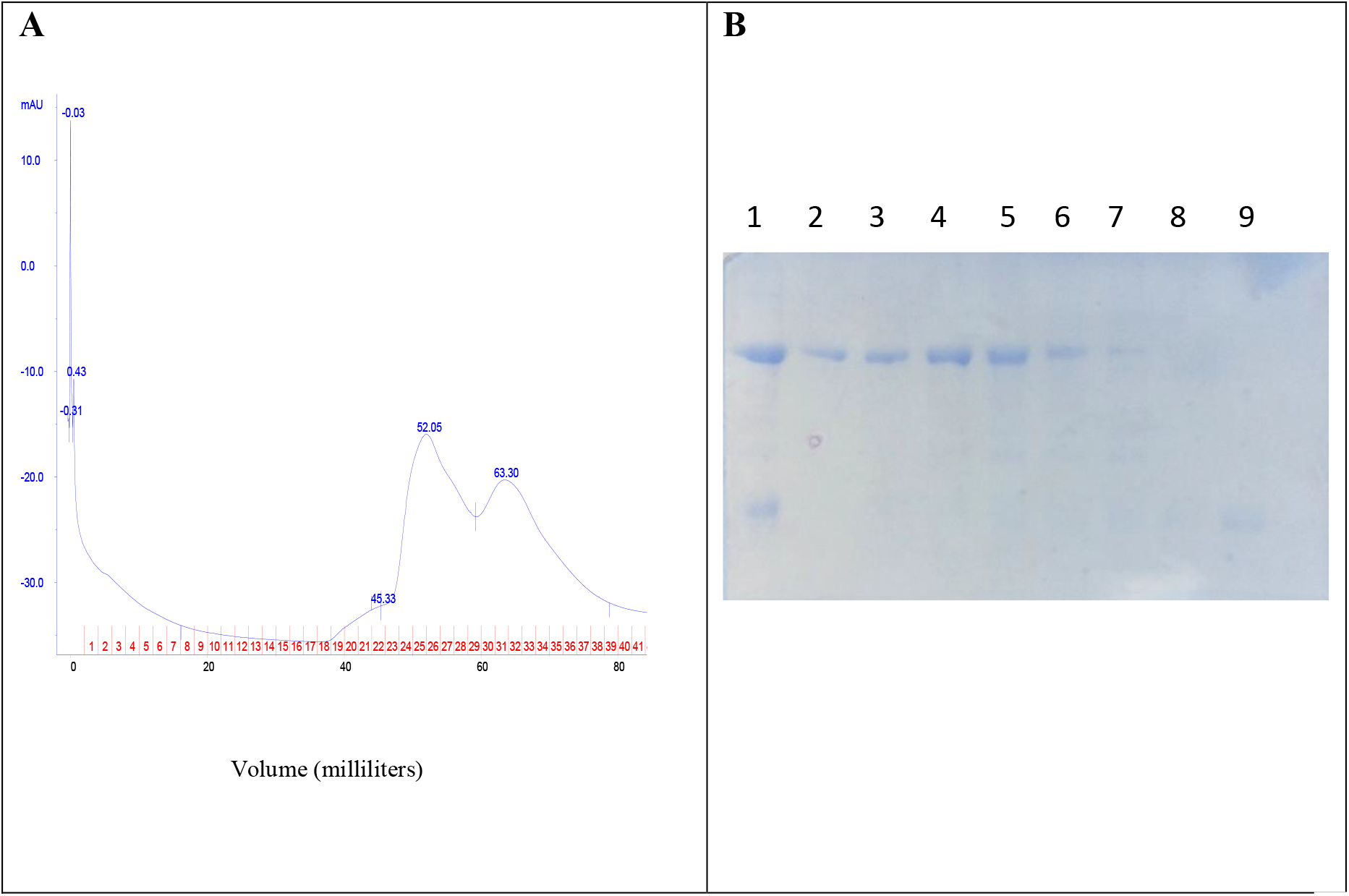
Size-exclusion Chromatography (Panel-A) and SDS-PAGE analysis (Panel-B) post heat- treated mixture of hFGF1 and cpSRP43. Panel-B: Lane 1= Preload to SEC containing cpSRP43 and hFGF1, Lanes 2-7= Peak 1 (fractions containing cpSRP43), Lanes 8 and 9= Peak 2 (fractions containing hFGF1).

## Conclusions

cpSRP43 is a vital component of the cpSRP pathway for its ability to interact with key players in this system and chaperone LHCPs through the stroma to the thylakoid membrane in an ATP-independent manner. LHCPs are apt to misfold and aggregate in aqueous environments due to their high hydrophobic content consequent to being integral membrane proteins. Certain reports have indicated that cpSRP43 exists in free form as a dimer in the stroma, as a potential intermediate, functioning in the formation of cpSRP as well as a disaggregase for adrift LHCP molecules (5,6). cpSRP43 has been shown to recognize and bind the hydrophilic L18 sequence, between TM2 and TM3 of LHCP (8). The conserved Tyrosine at position 204 in ANK3 of cpSRP43 recognizes the FDPLGL sequence in L18 (7, 8, 15, 20, 22). However, this interaction alone would not be adequate to guard the trans-membrane domains (TMDs) from aggregation (47). Investigation into how cpSRP43 can protect the TMDs of LHCP has only recently been communicated. Mapping of interaction sites between cpSRP43 and LHCb5 was done by alkylation efficiency of LHCb5 cysteine mutants using N-ethylmaleimide followed by site-specific cross-linking and the results of this combination of experiments revealed that cpSRP43 makes contacts and protects all three TMDs of the LHCb5 substrate (47). In site-directed mutagenesis studies, residues were categorized into two classes being mutations which disrupted cpSRP43’s chaperoning of LHCP but not the binding of L18 and mutations in which both were disrupted (47). These experiments were the basis for designing a model of the interaction surface on cpSRP43 for the protection of the TMDs of LHCP which is comprised of hydrophobic surfaces on ANK4, the bridging helix at the N-terminal of the ankyrin regions and, the beta- hairpins along the ankyrin regions (47). Since cpSRP43 is a known chaperone of LHCPs, it is conceivable that the binding interface between these two proteins can be elucidated in detail upon adequate investigation. The binding interface between cpSRP43 and other substrates is likely similar. However, considering the composite structure of cpSRP43 and the intricate rearrangements which it undergoes to bind its known partners, defining the exact binding interface for cpSRP43 and the substrates in this study would require extensive examination. The binding interface proposed by McAvoy et al. is primarily made up of hydrophobic residues suggesting that the protection mode of cpSRP43 is driven by the formation of hydrophobic interactions. This theory is upheld in another recent study in which cpSRP43 has been identified as a chaperone for glutamyl-tRNA reductase (GluTR), the rate-limiting enzyme in tetrapyrrole biogenesis (49). Tetrapyrrole is at the core of chlorophyll and the biogenesis of chlorophyll is coordinated with that of LHCPs in order for the proper assembly of light-harvesting complexes to proceed (49). Here they found that a deficit of cpSRP43 as well as GluTR-binding protein (GBP) significantly reduced production of GluTR and lead to a reduction in chlorophyll biogenesis which suggests that both cpSRP43 and GBP play distinct roles in the stabilization of GluTR (35). This group was able to form a ternary complex between cpSRP43, L18 and GluTR allowing them to speculate that both substrates are not competing for the same binding surface on cpSRP43 (49). Upon further examination, cpSRP43 was found to interact with the N-terminal of GluTR (49). The N- and C-terminal domains of GluTR can bind to different regulators to issue various control mechanisms for the enzyme (50) GBP was already known to interact with a heme-binding domain on the N-terminal of GluTR and this group conducted bimolecular fluorescence complementation assays (BiFC) and HIS pull-down assays in order to determine which GluTR domain interacts with cpSRP43 (49). cpSRP43 was found to bind to GluTR-N64–163 which contains the heme-binding domain as well as part of a catalytic domain, adjacent to the heme-binding domain, in the N-terminal (49). GluTR, in excess, is prone to aggregation and by using an aggregation-predicting algorithm, TANGO(51), the catalytic domain of GluTR was predicted to be an aggregation-prone region (49). Although this is a newly discovered role for cpSRP43, it is not surprising that cpSRP43 is providing a chaperoning function for another protein within the chloroplast.

We have already seen that cpSRP43 undergoes differing conformational rearrangements upon binding to partners along the cpSRP pathway (52). In addition, the exceptional ability of cpSRP43 to solubilize a substrate is key for its chaperoning function and can be achieved through contacts being hydrophobic and electrostatic in essence (1, 4, 6, 8, 47, 53, 54). These exceptional properties of cpSRP43 may make it challenging to define the binding interface between cpSRP43 and other substrates outside of the cpSRP system. As mentioned earlier, cpSRP43 is extremely sensitive to point mutations wherein the binding of L18 and chaperoning effect of cpSRP43 on LHCP was lost (47). These results suggest that cpSRP43 has many key residues which make it able to specifically chaperone LHCP, however the data collected in this chapter has demonstrated that cpSRP43 can act as a generic chaperone. There are many factors which can trigger protein aggregation including changes in ionic strength, pH, temperature and any other factor which can lead to interfacial instabilities (48). In this study we focused primarily on the role of cpSRP43 in the inhibition of heat-induced aggregation. The chaperoning function of cpSRP43 in the cpSRP pathway resembles that of the Hsp104/ClpB family of proteins. Hsp104 and ClpB work in combination with Hsp70 or DnaK to disaggregate and remodel proteins under severe stress conditions and requires energy from ATP to perform (20). Although this family of proteins are impressive disaggregating machines, it is noteworthy to consider that cpSRP43 has the same thermoresistance and, in addition, can disaggregate and remodel the LHCP molecules without the input of external energy. Here we have shown that cpSRP43 has the capability to delay or inhibit the heat-induced aggregation of proteins of diverse sizes and characteristics. CA, a 30 kDa enzyme, was able to resist aggregation in the presence of cpSRP43 upon the application of heat and subsequent to denaturation and reduction. Con A, a homotetramer with each subunit having a molecular weight of 26.5 kDa showed substantial reduction in its propensity to aggregate in the presence of cpSRP43 as compared to the samples of Con A which were incrementally heated in the absence of cpSRP43. The rate of aggregation of hFGF1 was significantly reduced while the bioactivity was retained after being heated in the presence of cpSRP43 as compared to in the absence of cpSRP43 wherein most of the hFGF1 became aggregated. The best protection in each scenario was found at ratios of substrate to cpSRP43 in which cpSRP43 was in higher abundance supporting the theory that this is a concentration-dependent function which would require regulation. Given the information known about the interaction between cpSRP43 and LHCP as well as the more recent report in which cpSRP43 is serving as a chaperone for GluTR, it is feasible to conclude that the mechanism of protection is mainly interchangeable between substrates with some slight differences. cpSRP43 can predominately form hydrophobic interactions with its substrate thereby shielding it from aggregation or aiding in protein refolding. Although we know that cpSRP43 is more successful when present in excess to its substrate, the question remains as to how many molecules of cpSRP43 are interacting along each target protein. This information is expected to vary depending on the substrate. In this study, the chaperoning effect of cpSRP43 has been extended to and substantiated by proteins which are not native to the chloroplast. In the case of all three candidate proteins requiring chaperone facilitation, either the existence of aggregation was not detected, or the rate of aggregate accumulation was significantly lowered in the presence of cpSRP43. The exact mechanism by which cpSRP43 can successfully provide protection to these proteins requires further investigation. It is highly anticipated that many more roles for cpSRP43 in the chloroplast will be revealed and now that we have displayed the ability of cpSRP43 to act as a generic chaperone, further investigation using a variety of substrates may lead to the use of cpSRP43 in applied biochemical and medical science.

## Acknowledgements

This project was supported by the Department of Energy (DE-FG02-01ER15161 and the Arkansas Biosciences Institute.

